# Model-based spike sorting with a mixture of drifting *t*-distributions

**DOI:** 10.1101/109850

**Authors:** Kevin Q. Shan, Evgueniy V. Lubenov, Athanassios G. Siapas

## Abstract

Chronic extracellular recordings are a powerful tool for systems neuroscience, but spike sorting remains a challenge. A common approach is to fit a generative model, such as a mixture of Gaussians, to the observed spike data. Even if non-parametric methods are used for spike sorting, such generative models provide a quantitative measure of unit isolation quality, which is crucial for subsequent interpretation of the sorted spike trains. We present a spike sorting strategy that models the data as a mixture of drifting *t*-distributions. This model captures two important features of chronic extracellular recordings—cluster drift over time and heavy tails in the distribution of spikes—and offers improved robustness to outliers. We evaluate this model on several thousand hours of chronic tetrode recordings and show that it fits the empirical data substantially better than a mixture of Gaussians. We also provide a software implementation that can re-fit long datasets (several hours, millions of spikes) in a few seconds, enabling interactive clustering of chronic recordings. Using experimental data, we identify three common failure modes of spike sorting methods that assume stationarity. We also characterize the limitations of several popular unit isolation metrics in the presence of empirically-observed variations in cluster size and scale. We find that the mixture of drifting *t*-distributions model enables efficient spike sorting of long datasets and provides an accurate measure of unit isolation quality over a wide range of conditions.

## 1. Introduction

Chronic extracellular recordings offer access to the spiking activity of neurons over the course of days or even months. However, the analysis of extracellular data requires a process known as spike sorting, in which extracellular spikes are detected and assigned to putative sources. Despite many decades of development, there is no universally-applicable spike sorting algorithm that performs best in all situations.

Approaches to spike sorting can be divided into two categories: model-based and non-model-based (or non-parametric). In the model-based approach, one constructs a generative model (e.g. a mixture of Gaussian distributions) that describes the probability distribution of spikes from each putative source. This model may be used for spike sorting by comparing the posterior probability that a spike was generated by each source. Fitting of such models may be partially or fully automated using maximum likelihood or Bayesian methods, and the model also provides an estimate of the misclassification error.

In the non-parametric approach, spike sorting is treated solely as a classification problem. These classification methods may range from manual cluster cutting to a variety of unsupervised learning algorithms. Regardless of the method used, scientific interpretation of the sorted spike train still requires reliable, quantitative measures of unit isolation quality. Often, these heuristics either explicitly (Hill et al., 2011) or implicitly (Schmitzer-Torbert et al., 2005) assume that the spike distribution follows a mixture of Gaussian distributions.

However, a mixture of Gaussians does not adequately model the cluster drift and heavy tails that are observed in experimental data (Figure 1). Cluster drift is a slow change in the shape and amplitude of recorded waveforms (Figure 1C), usually ascribed to motion of the recording electrodes relative to the neurons. This effect may be small for short recordings (< 1 hour), but can produce substantial errors if not addressed in longer recordings (Figure 5). Spike clusters also have heavier tails than expected from a Gaussian distribution (Shoham et al., 2003), and may be better fit using a multivariate *t*-distribution (Figure 1D and Figure 4).

**Figure 1.**
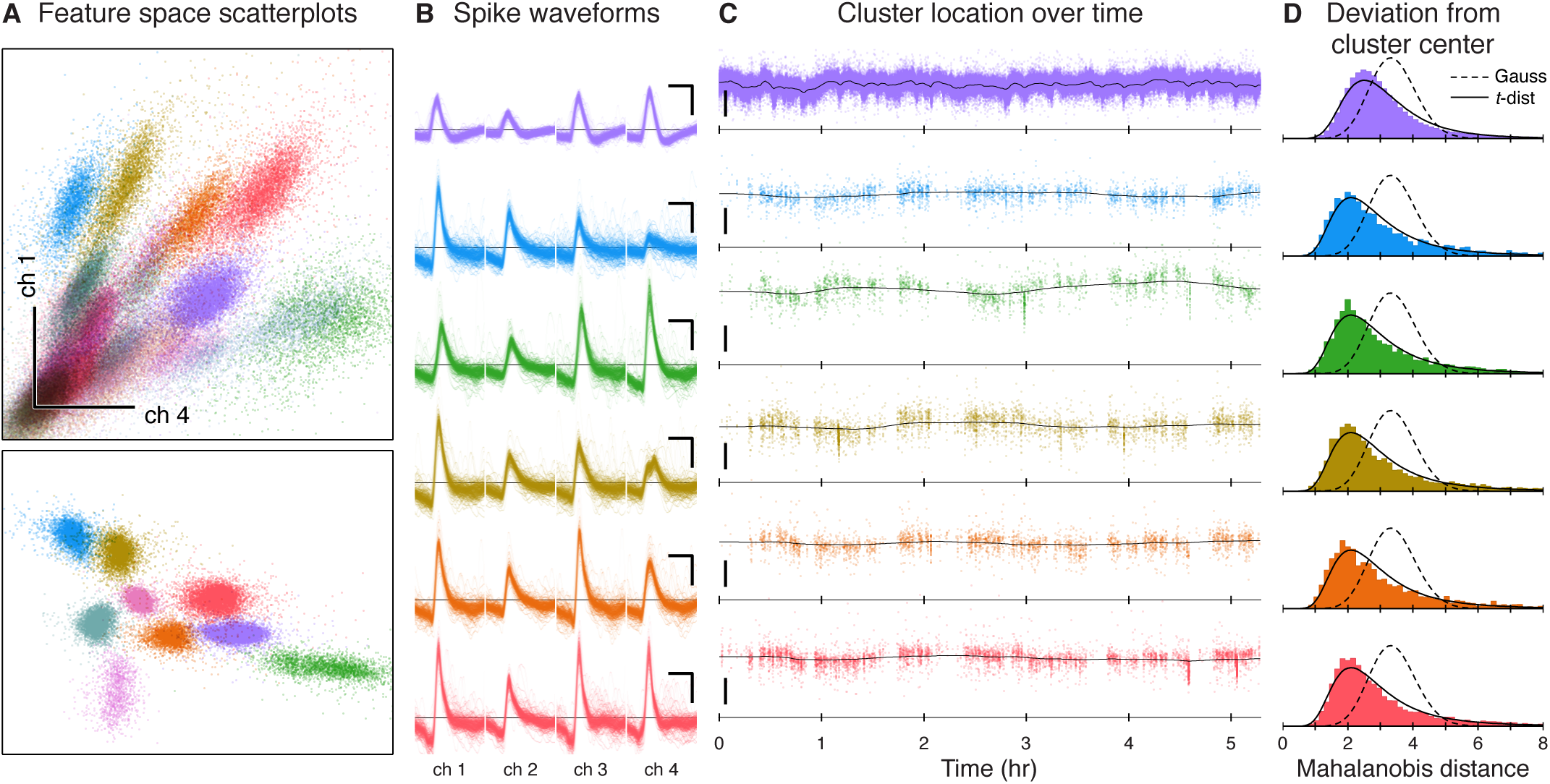
Extracellular recordings contain drifting, heavy-tailed clusters. (**A**) Scatterplots of spikes in feature space, color-coded by putative identity. Spike waveforms recorded on 4 tetrode channels were projected onto a 12-dimensional feature space using 3 principal components from each channel. Top: scatterplot of the first principal component from channels 1 and 4. Bottom: a different projection of the data, showing only the best-isolated single units. Scale bar: 50 μV RMS. (**B**) Spike waveforms (inverted polarity) for six example units. Scale bar: 200 μV, 0.5 ms. (**C**) Cluster drift over time. Black line indicates the cluster center fitted using the MoDT model. Scale bar: 50 μV RMS. (**D**) Observed and theoretical distributions of the Mahalanobis distance from the fitted cluster center to the assigned spikes. If the spikes were Gaussian-distributed, the squared distance should follow a chi-squared distribution (dashed line). Instead, the observed distances follow the theoretical distribution for *t*-distributed spikes (solid line; see Appendix C.1 for derivation).

**Figure 4.**
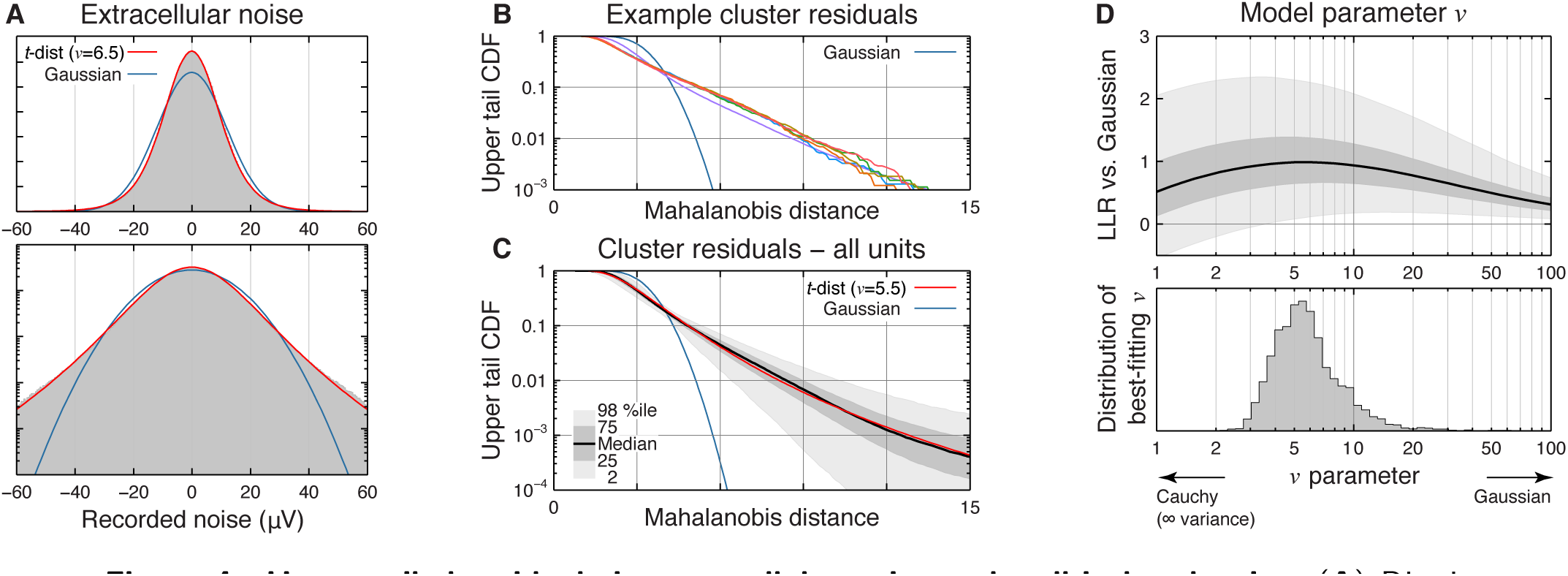
Heavy-tailed residuals in extracellular noise and well-isolated units. (**A**) Distribution of extracellular noise for an example tetrode channel during periods where no spikes were detected. Red and blue lines show a *t*-distribution and Gaussian fit, respectively. Bottom panel shows the same histogram with a logarithmic y-scale. (**B**) Upper tail CDF (fraction of a unit’s spikes that lie beyond a given Mahalanobis distance from the cluster center) for the 6 example units from Figure 1. (**C**) Upper tail CDF for all well-isolated units. Shaded areas indicate quantiles across units. Theoretical distributions for a *t*-distribution and Gaussian are shown for reference. (**D**) Effect of changing the model’s *v* parameter, which controls the heavy-tailedness of the distribution. Top: Log-likelihood ratio (LLR per spike) comparing the *t*-distribution vs. Gaussian model for all units. Bottom: Overall distribution of best-fitting for *v* each unit.

**Figure 5.**
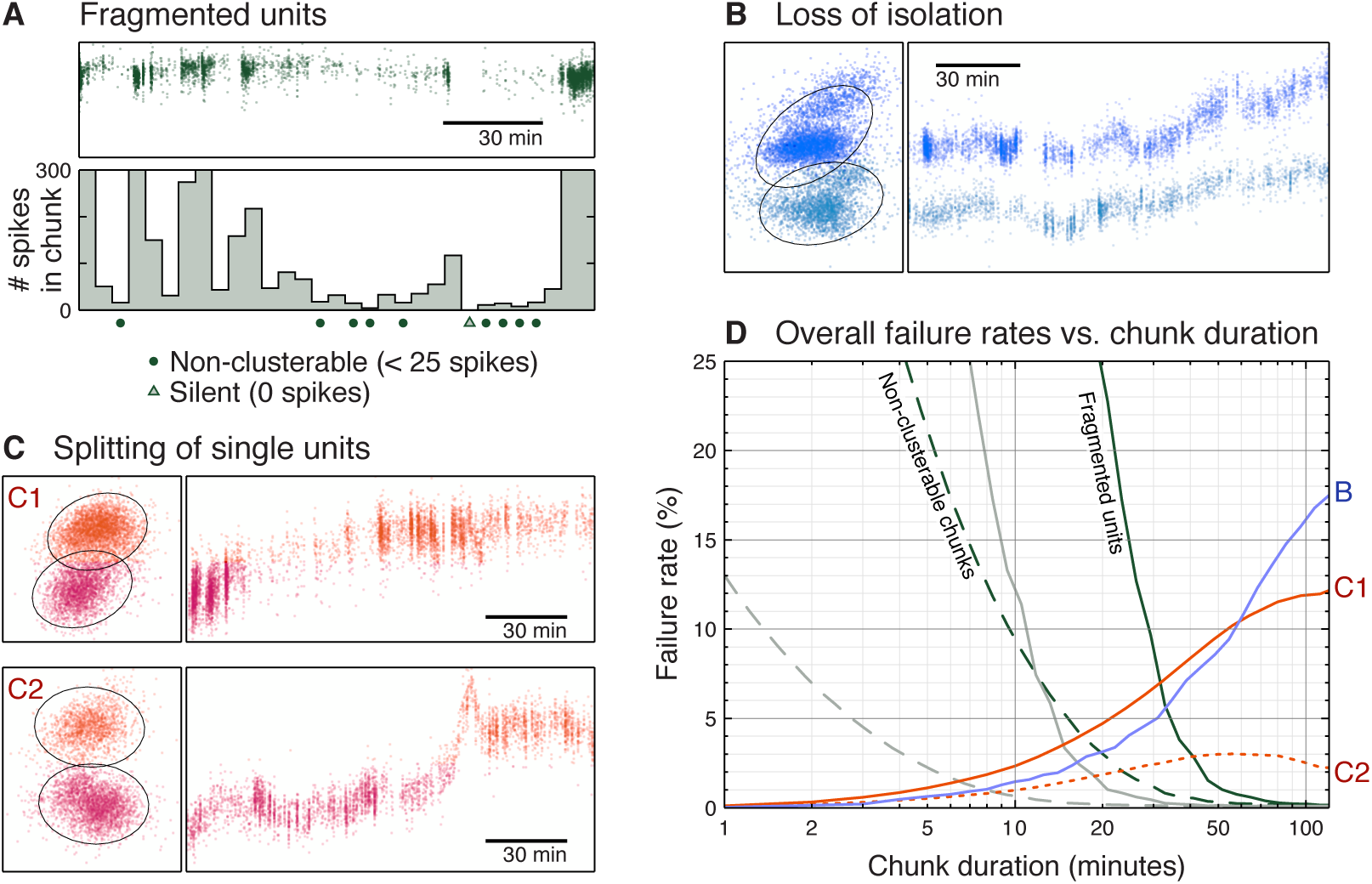
Failure modes of a stationary approach. An alternative approach to handling cluster drift is to break the recording into chunks, perform spike sorting on each chunk independently, and finally link the spike clusters over time. We identified three common failure modes of this approach. (**A**) In order to link a unit over time, each chunk must contain enough of that unit’s spikes to form a cluster (we used a threshold of 25 spikes). If any chunk is non-clusterable, then the unit cannot be successfully linked and will become fragmented. Panel D shows the overall prevalence of these failures for varying chunk durations (dashed and solid green lines). The light grey lines (dashed and solid) repeat this analysis with the clusterability threshold set at 1 spike. (**B**) Drift causes clusters to become smeared out over time. If we analyze these irregularly-shaped clusters as stationary distributions, then some units will appear to overlap even though they remain well-isolated over time. This loss of isolation artificially reduces the yield of good units. (**C**) Drift may produce a multi-modal density distribution. As a result, the clustering algorithm may split a single unit into two clusters (C1). In some cases, these two clusters may be quite well-isolated from each other (C2). (**D**) Overall prevalence of these failure modes for varying chunk durations.

To address these issues, we model the spike data as a mixture of drifting t-distributions (MoDT). This model builds upon previous work that separately addressed the issues of cluster drift (Calabrese & Paninski, 2011) and heavy tails (Shoham et al., 2003), but we have found the combination to be extremely powerful for modeling and analyzing experimental data. We also discuss the model’s robustness to outliers and describe some algorithmic modifications that reduce the computational runtime.

We used the MoDT model to perform spike sorting on 34,850 tetrode-hours of chronic tetrode recordings (4.3 billion spikes) from the rat hippocampus, cortex, and cerebellum. Using these experimental data, we evaluate the assumptions of our model and provide recommended values for the model’s user-defined parameters. We also analyze how the observed cluster drift may impact the performance of spike sorting methods that assume stationarity. Finally, we investigate how the observed variations in cluster size, scale, and tail distribution may impact the reliability of several unit isolation metrics.

## 2. Methods

### 2.1. Mixture of drifting *t*-distributions (MoDT) model

Spike sorting begins with spike detection and feature extraction. During these preprocessing steps, spikes are detected as discrete events in the extracellular voltage trace and represented as points *y_n_* in some *D*-dimensional feature space.

The standard mixture of Gaussians (MoG) model treats this spike data *y*_n_ as samples drawn from a mixture distribution with PDF given by

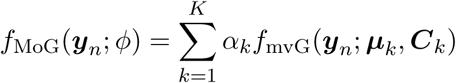

where *ϕ* = {…, *α_k_*, *μ_k_*, *C_k_*,…} is the set of fitted parameters, *K* is the number of mixture components, *α_k_* are the mixing proportions, and *f*_mvG_(*y; μ, C*) is the PDF of the multivariate Gaussian distribution with mean *μ* and covariance *C*:

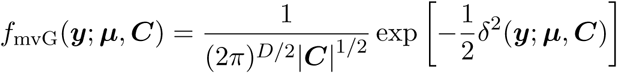

For notational convenience, let *δ*^2^ denote the squared Mahalanobis distance

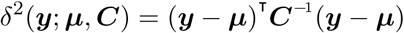

We make two changes to this model. First, we replace the multivariate Gaussian distribution with the multivariate *t*-distribution. The PDF for this distribution, parameterized by location *μ*, scale *C*, and degrees-of-freedom *v*, is given by

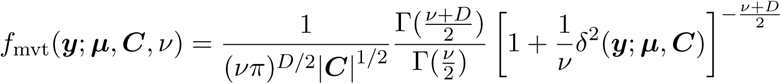

Second, we break up the dataset into *T* time frames (we used a frame duration of 1 minute) and allow the cluster location *μ* to change over time. The mixture distribution becomes

**Table 1.** Mathematical notation. Lowercase bold letters (*y_n_*, *μ_kt_*) denote *D*-dimensional vectors, and uppercase bold letters (*C_k_*, *Q*) denote *D* × *D* symmetric positive definite matrices.

|  |  |
| --- | --- |
| Dimensions |  |
| $D$ | Number of dimensions |
| $N$ | Number of spikes |
| $K$ | Number of clusters |
| $T$ | Number of time frames |
| Given data |  |
| $\mathbf{y}_n$ | Observed spike $n$ |
| $t_n$ | Time frame in which spike $n$ occurred |
| $w_n$ | Weighting of spike $n$ (multiplier applied to log-likelihood) |
| User-defined constants |  |
| $\nu$ | $t$-distribution degrees-of-freedom parameter |
| $\mathbf{Q}$ | Drift regularization parameter |
| Fitted model parameters |  |
| $\alpha_k$ | Mixing proportion for cluster $k$ |
| $\boldsymbol{\mu}_{kt}$ | Location parameter for cluster $k$ in time frame $t$ |
| $\mathbf{C}_k$ | Scale parameter for cluster $k$ |
| Latent variables introduced by EM procedure |  |
| $z_{nk}$ | Posterior probability that spike $n$ belongs to cluster $k$ |
| $u_{nk}$ | Scaling variable introduced in formulating the $t$ distribution as a Gaussian-Gamma compound distribution |

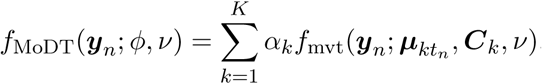

where *t_n_* ∈ {1,…,*T*} denotes the time frame for spike *n*. We use a common *v* parameter for all components and have chosen to treat it as a user-defined constant. The fitted parameter set is thus *ϕ* = {…, *α_k_*, *μ*_*k*1_,…, *μ_kT_*, *C_k_*,…}.

In order to enforce consistency of the component locations across time, we introduce a prior on the location parameter that penalizes large changes over consecutive time steps. This prior has a joint PDF proportional to

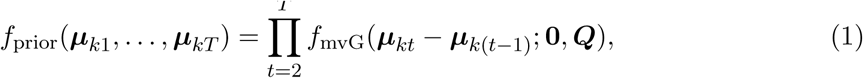

where *Q* is a user-defined covariance matrix that controls how much the clusters are expected to drift.

### 2.2. EM algorithm for model fitting

Assuming independent spikes and a uniform prior on the other model parameters, we can obtain the maximum *a posteriori* (MAP) estimate of the fitted parameters *ϕ* by maximizing the log-posterior, which is equivalent (up to an additive constant) to the following:

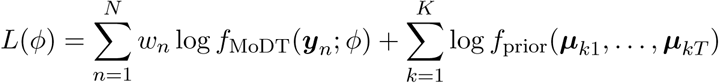

Note that we have introduced a weight *w_n_* for each spike. This allows us to fit the model to a subset of the data while remaining consistent with the full dataset (Feldman et al., 2011).

As with most mixture distributions, it is intractable to optimize *L*(*ϕ*) directly. However, by introducing additional latent random variables, we obtain a “complete-data” log-posterior *L_c_*(*ϕ, Z, U*) that allows us to decompose the problem and optimize it using an expectation-maximization (EM) algorithm (McLachlan & Peel, 2000).

In the E-step, we compute the expected value of *L_c_* assuming that these latent variables follow their conditional distribution given the observed data and the fitted parameters 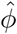 from the previous EM iteration. The conditional expectations of these latent variables are given by:

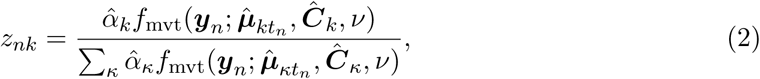

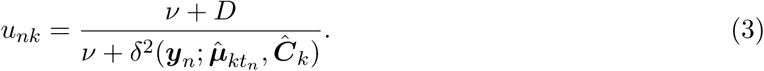

The *z*_*nk*_ correspond to the posterior probability that spike *n* was produced by component *k*, and may thus be used for spike sorting. The *u_nk_* arises from the formulation of the *t*-distribution as a Gaussian-Gamma compound distribution and may be interpreted as a scaling variable that “Gaussianizes” the multivariate *t*-distribution. In the Gaussian case (the limit of a *t*-distribution as *v* ⟶ ∞), we have *u*_*nk*_ = 1 for all spikes. For finite *v*, note that *u*_*nk*_ decreases as the Mahalanobis distance *δ* increases.

Next we can compute the conditional expectation of *L_c_*(*ϕ, Z, U*) over these latent variables. Following Peel & McLachlan (2000), we find that this is equivalent (up to an additive constant) to

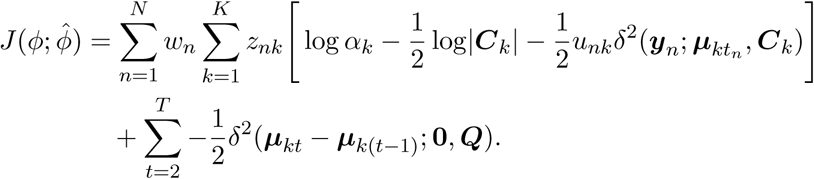

In the M-step, we maximize
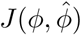
 with respect to the fitted parameters. The optimal value for the mixing proportions *α* is simply a weighted version of the mixture of Gaussians (MoG) M-step update:

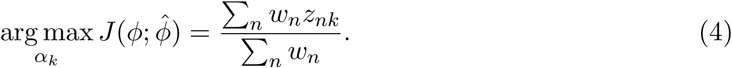

The optimal value for the cluster scale parameter ***C*** is also similar to the MoG case, but each spike is additionally scaled by *u_nk_*:

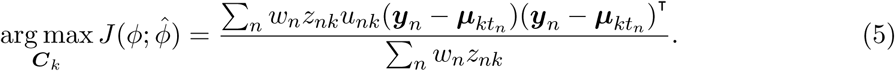

For the cluster location parameters *μ*, note that 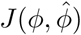
 is quadratic with respect to *μ* and its maximum occurs where the gradient 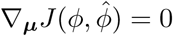
. Hence we can find the optimal *μ* by solving the following linear system of equations:

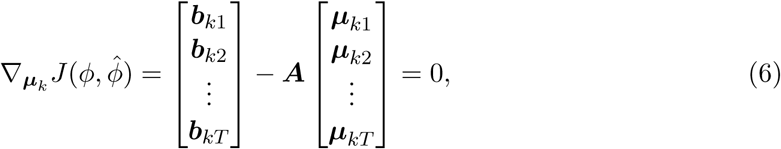

where

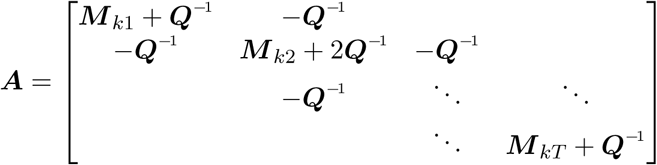

and

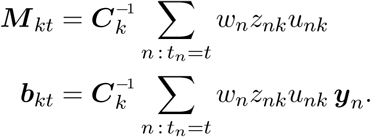

Although this is a *DT* × *DT* linear system, its sparsity structure allows us to solve for ***μ*** with a complexity that scales linearly with *T* (Paninski et al., 2010). When *T* ≪ *N*, solving equation (6) accounts for a negligible fraction of the overall computational runtime. Appendix A describes some alternative methods for the M-step update of ***μ***.

Note that the scaling variable *u_nk_* acts as an additional weighting term in the optimization of ***μ*** and ***C***. Since *u_nk_* decreases as spike *n* gets far away from cluster *k*, any outliers are automatically discounted during the fitting process. As a result, the fitted parameters are considerably more robust to the presence of outliers than in the Gaussian case (Figure 2). This property makes the *t*-distribution a useful model even when the underlying data are Gaussian.

**Figure 2.**
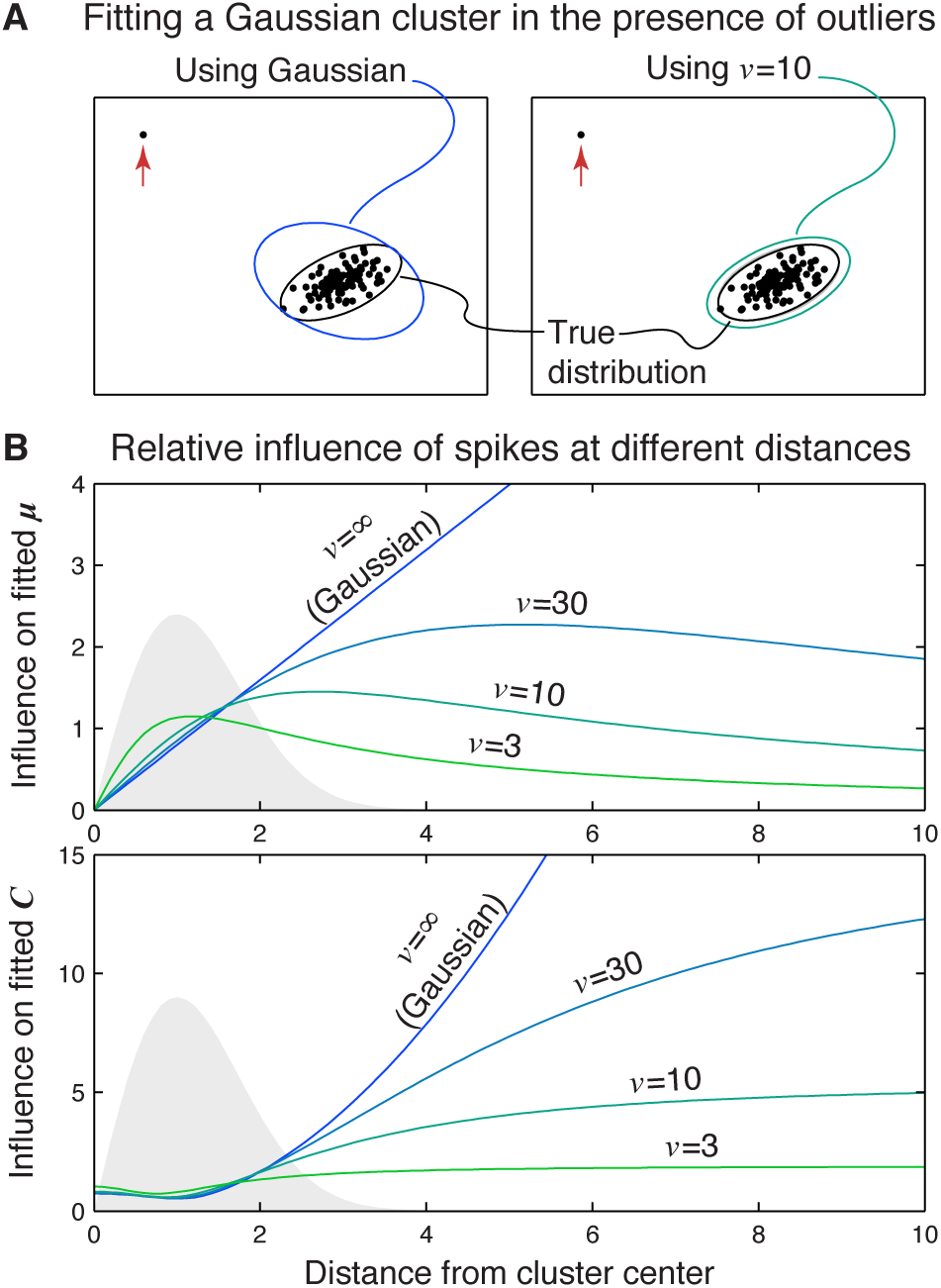
Fitted parameters of the *t*-distribution are robust to outliers. (**A**) In this example we generated 100 points from a Gaussian distribution and added a single outlier (red arrow). This has stretched out the estimated covariance when using a Gaussian fit (left). In contrast, fitting with a *t*-distribution (right) comes much closer to the true parameters. (**B**) For a Gaussian model, the relative influence of a single spike grows unbounded with increasing distance from the cluster center, allowing outliers to exert an undue influence on the fitted model parameters. When fitting a *t*-distribution, the scaling variable *u*_*nk*_ effectively discounts any spikes far away from the cluster center, thereby limiting the effect of outliers. The grey histogram in the panel background shows the theoretical distance distribution for a Gaussian cluster. See appendix C.3 for more detail.

We provide a MATLAB implementation of this EM algorithm that may be downloaded from https://github.com/kqshan/MoDT. It offers a mild speedup over the MATLAB built-in mixture of Gaussians fitting routine, despite fitting a more complex model (Table 2). In addition, it supports the use of a weighted training subset, which offers a proportional reduction in runtime at the expense of model accuracy. It also supports the use of GPU computing using the NVidia CUDA computing platform, which offers an additional 3-20x speedup, depending on the hardware.

**Table 2.** Computational runtime. Time required to perform 20 EM iterations on the sample dataset shown in Figure 1 (D=12, K=26, N=1.9 million). fitgmdist is a mixture of Gaussians fitting routine that is part of the MATLAB Statistics and Machine Learning Tool-box. modt is a MATLAB implementation of our EM algorithm that may be downloaded from https://github.com/kqshan/MoDT. In addition to fitting the richer MoDT model, it supports two additional features (data weights and GPU computing) that can dramatically reduce fitting times. Benchmarking was performed on an Amazon EC2 g2.2xlarge instance running MATLAB R2016a.

| Model type | Algorithm description | Runtime (s) |
| --- | --- | --- |
| MoG | `fitgmdist` | 208.21 |
| MoG | `modt` in Gaussian mode | 178.30 |
| MoDT | `modt` | 186.76 |
| MoDT | `modt` on GPU | 22.93 |
| MoDT | `modt`, 5% subset | 9.83 |
| MoDT | `modt`, 5% subset, GPU | 2.71 |

### 2.3. Interactive clustering

We relied on human operators to guide the model fitting in an interactive clustering step. The user can change the number of clusters (*K*) by choosing clusters to split or merge, and we re-fit the model after each operation. To ensure an adequate user experience, we used weighted training subsets as necessary to ensure that we could re-fit the model and update the user interface in a matter of seconds.

The quality of the initial fit is an important factor in determining the user workload. When the dataset comes from a sequence of recordings with stable electrodes, we initialize the model using the fitted parameters from the previous dataset. Using this initialization, we have found that most datasets require minimal user interaction.

When no prior dataset exists, we use a split-and-merge technique (Ueda et al., 2000) for parameter initialization and the Bayes information criterion (BIC) for model selection, similar to the method described by Tolias et al. (2007). This initialization works well in brain areas with a low density of neurons (e.g. cortex), but typically requires manual intervention in brain areas with greater multi-unit activity, especially if the units of interest fire very sparsely (e.g. hippocampal area CA1).

### 2.4. Data preprocessing

To evaluate the MoDT model for spike sorting, we generated 34,850 tetrode-hours (34 terabytes) of chronic tetrode data by implanting 10 Long-Evans rats with 24-tetrode arrays targeting areas of the hippocampus, cortex, and cerebellum. Extracellular signals were digitized at 25 kHz and recorded continuously, but these recordings were broken into datasets ranging from 1 to 25 hours in duration, depending on experimental needs. All animal procedures were in accordance with National Institutes of Health (NIH) guide for the care and use of laboratory animals, and approved by the Caltech Institutional Animal Care and Use Committee.

We detected spikes offline by bandpass filtering the data using an FIR filter with a 600-6,000 Hz passband, upsampling to 50 kHz (Blanche & Swindale, 2006), and identifying peaks in the nonlinear energy operator (Mukhopadhyay & Ray, 1998). Overall, we detected and sorted 4.3 billion spikes.

We performed dimensionality reduction on the spike waveforms by projecting them onto a set of basis waveforms. These were chosen using principal component analysis (PCA) to select 3 PCA axes from each of the 4 tetrode channels (12 dimensions in total). We used a robust version of principal components analysis (using the Huber loss function; see e.g. Udell et al., 2016) to ensure that the chosen axes would not be unduly influenced by high-amplitude artifacts.

This dimensionality reduction preserves the root-mean-square (RMS) amplitude of the spike waveforms along the PCA axes. Although these RMS amplitudes have units of microvolts, these values are smaller than the waveform peak amplitudes that are commonly reported in spike sorting. For example, the blue unit in Figure 1 (second row in panels B-D) has an RMS amplitude of 100 μV along the first PCA feature dimension on channel 1, but its peak amplitude on that channel is 384 μV.

If two neurons fire near-simultaneously, their spikes will overlap and produce a waveform that is the sum of both waveforms. If both spikes are detected, we can mitigate this distortion by deconvolving the overlapped waveforms during the dimensionality reduction step. However, many overlapped waveforms are detected as a single spike, and these appear as outliers during the clustering process. Thanks to the *t*-distribution’s robustness to outliers, we can ignore these spikes during clustering, and then identify and reassign them in a post-processing step. If necessary, template matching may be performed using template waveforms derived from the fitted cluster locations (Prentice et al., 2011; Pillow et al., 2013).

### 2.5. Computational infrastructure

All analysis was implemented in MATLAB and performed on a small computer cluster consisting of 4 compute nodes (a total of 64 cores) and 11 storage nodes (serving data using the NFS network file system). For non-interactive tasks, we used the MATLAB Parallel Computing Toolbox to submit batch jobs to a Sun Grid Engine job scheduler. We used the DataJoint MATLAB toolbox (Yatsenko et al., 2015) with a MySQL relational database to (1) keep track of which datasets were waiting in the compute queue and which were ready for interactive clustering, (2) automate the computation of derived quantities and (3) store all metadata, ranging from recording parameters to unit quality metrics, in an efficiently-queryable manner.

The interactive clustering step was the bottleneck of the spike sorting process due to the small number of human users (4 users vs. 64 compute cores) and their low availability (only a few hours per day, compared to the 99% uptime of the computer cluster). We did not explicitly track the amount of time spent clustering, but based on file modification timestamps, we estimate that it took 500 user-hours to cluster these 34,850 tetrode-hours of data. Average throughput ranged from 13-27 datasets per user-hour, independent of the dataset’s recording duration.

Benchmarking (Table 2) was performed on an Amazon Elastic Compute Cloud (EC2) g2.2xlarge instance running MATLAB R2016a (64-bit) on Ubuntu 14.04.4 LTS with CUDA toolkit 7.5. This virtual machine has access to 8 threads of an Intel Xeon E5-2670 CPU, 15 GB of memory, and one GPU of an NVidia Grid K520 graphics acceleration board. All floating-point computations were performed in double precision.

### 2.6. Measuring unit isolation using the MoDT model

The MoDT model may be fitted to previously spike-sorted data by substituting the given spike assignments for the latent variable *z_nk_*

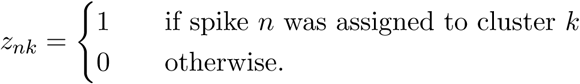

If the spike sorting algorithm can provide soft assignments (i.e. each spike has a probability of belonging to each cluster rather than being fully assigned to a single cluster), then these may be used instead. With *z_nk_* fixed, model fitting typically requires fewer than 10 EM iterations.

After fitting, (equation (2) provides a model-based estimate of the probability that spike *y_n_* was produced by each of the source clusters. Summing these *z_nk_* provides the expected number of misclassified spikes. Following Hill et al. (2011), we define the false positive (FP) fraction and the false negative (FN) ratio for cluster *k* as

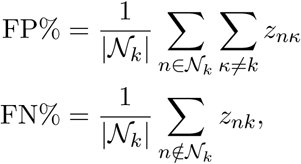

where *N_k_* is the set of spikes assigned to cluster *k*. In the hypothesis testing literature, the FP fraction is also known as the false discovery rate. The FN ratio does not have a similar analogue, and it may be greater than one.

The MoDT model also provides a natural generalization of Gaussian-based unit isolation metrics to the drifting case. By setting *v* = ∞, the fitted ***μ**_kt_* and ***C**_k_* correspond to the time-varying cluster mean and the cluster covariance, respectively. These were used to compute Mahalanobis distances for the empirical comparison of unit isolation quality metrics (Figure 6E).

**Figure 6.**
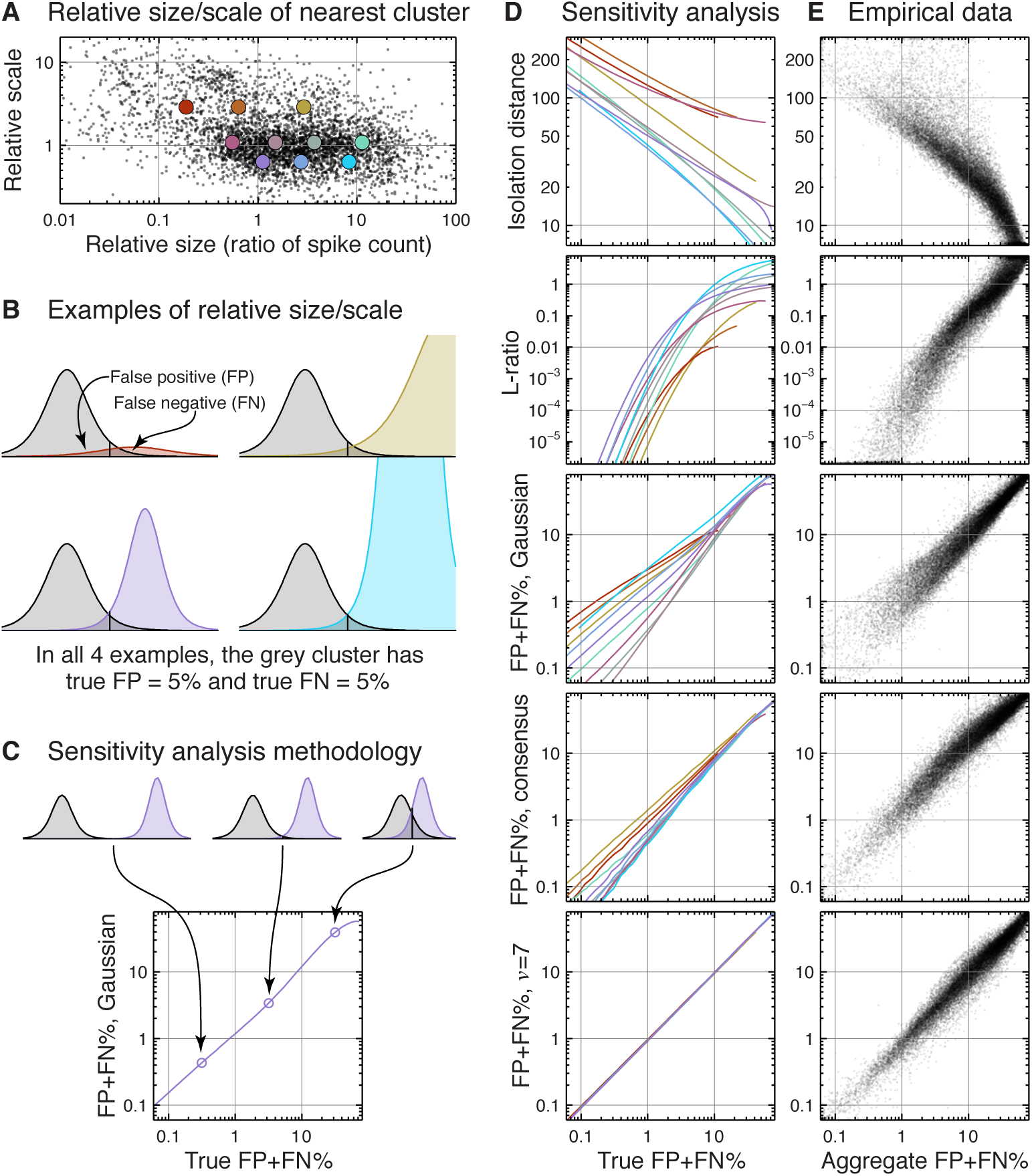
Sensitivity analysis of unit isolation metrics. (**A**) Relative size and scale of the nearest neighboring cluster for each of our 4,432 well-isolated units. This shows that potential sources of contamination come in a wide variety of sizes and scales. We selected 10 representative cases (colored dots) for our sensitivity analysis. (**B**) We performed our sensitivity analysis by generating synthetic clusters and comparing our unit isolation metrics to the true misclassification error. These four example cases all have true false positive (FP) fraction of 5% and a false negative (FN) ratio of 5%, but use a different size/scale for the contaminating cluster. (**C**) As we move the contaminating cluster closer or farther away, we trace out a curve relating the unit isolation metric to the true error. We repeat this process for each of the 10 cases shown in panel A. (**D**) Sensitivity analysis results. Ideally, each of the 10 colored curves should lie on top of one another, indicating that the metric is not sensitive to these variations in cluster size and scale. (**E**) Comparison of isolation metrics on experimental data. Since the true error is unknown, these metrics are compared to an aggregate error metric that is based on all five metrics combined.

### 2.7. Unit selection criteria

This spike sorting process identified 20,630 putative single units, accounting for 852 million spikes over 89,127 unit-hours.

However, to mitigate the risk of our analysis results being confounded by inaccurate spike sorting, we applied a more conservative set of selection criteria than usual. We required that units have a low misclassification error (estimated false positive and false negative each less than 2%), be reliably detected (waveform peak amplitude greater than 200 μV and estimated spike detection false negative rate less than 2%), exhibit a clean refractory period, and come from a recording at least 2 hours in duration.

Using these criteria, we identified 4,432 very well-isolated single units, accounting for 338 million spikes over 32,890 unit-hours. The median quality metrics of these units are as follows: 0.5% false positive and 0.8% false negative due to misclassification, 340 μV waveform peak amplitude, 0.0005% false negative from the spike detection threshold, and 0.4% false positive based on refractory period violations (see Hill et al., 2011, for the definition of these error rates).

## 3. Results

We analyze these 4,432 well-isolated single units in this section. First, we quantify the cluster drift and the distribution of spike residuals to provide recommendations for user-defined parameters in our model. Next, we use these units—which we have tracked over several hours at a time—to consider the consequences of using a stationary model for spike sorting. Finally, we evaluate how some commonly-used unit isolation metrics are affected by the heavy tails and other features encountered in these experimental data.

### 3.1. Cluster drift in empirical data

We quantified the cluster drift by measuring the distance from each unit’s current location (determined using a 40-minute moving average) to its location at the start of the recording. Individual clusters may move closer or farther away from where they started (Figure 3A), but over all units, the average distance increases and the distribution spreads out (Figure 3B). These distances are measured in feature space units, which correspond to root-mean-square (RMS) waveform amplitude in microvolts.

**Figure 3.**
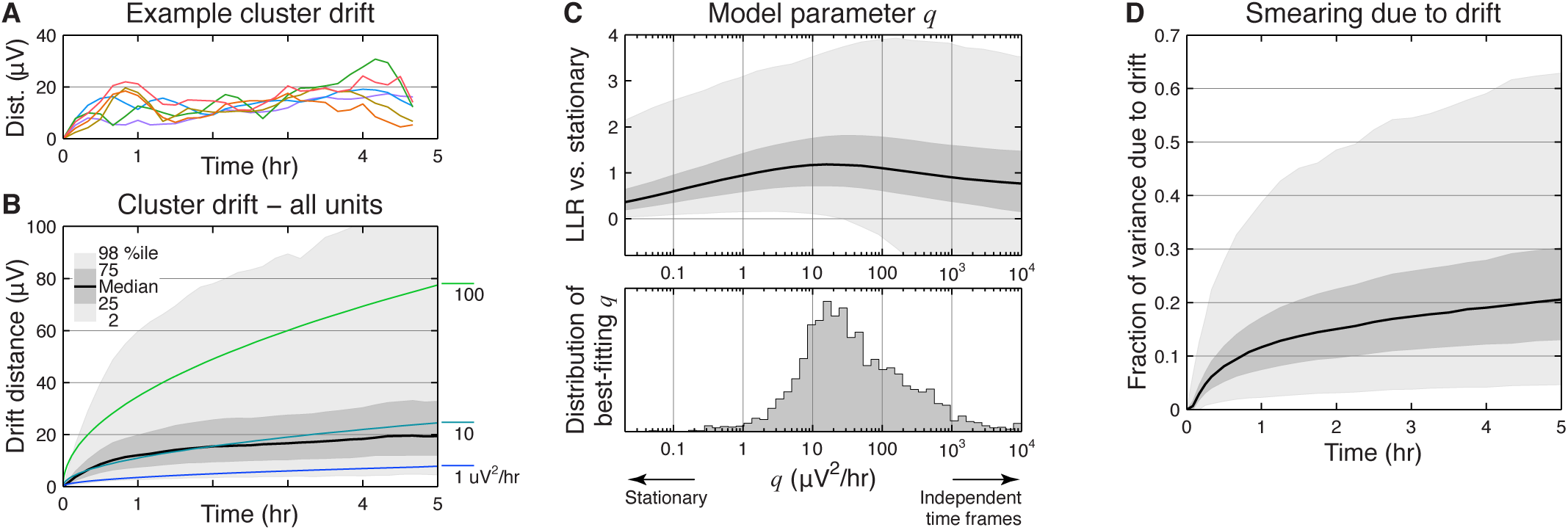
Cluster drift in well-isolated units. (**A**) Cluster drift of the 6 example units from Figure 1. Distances are measured from the unit’s current location (determined using a 40-minute moving average) to its location at the start of the dataset. (**B**) Cluster drift of all well-isolated units in our analysis. Shaded regions indicate quantiles across units. Colored lines show expected drift distances for 3 different drift rates. (**C**) Effect of changing the model’s drift regularization parameter **Q**. We only considered isotropic matrices ***Q*** = *q**I***. Top: Test-set log-likelihood ratio (LLR per spike) comparing the drifting vs. stationary model for all units. Bottom: Overall distribution of the best-fitting *q* for each unit. (**D**) If drift is not accounted for, it produces an apparent “smearing” of the spike distribution in feature space. This panel shows the fraction of spike variability that can be accounted for by cluster drift.

Note that the cluster location prior in equation (1) corresponds to a Gaussian random walk with a constant rate of drift. However, the observed distribution of drift distances is much broader than expected from such a process, and so this aspect of the MoDT model should be treated as a regularizer rather than an attempt to accurately model the underlying phenomena.

The MoDT model parameter ***Q*** is a user-defined constant that controls this regularization. Figure 3C shows the effect of changing this parameter over a wide range of values. The log-likelihood ratio (LLR) is a measure of the MoDT model’s quality of fit compared to a stationary alternative; values greater than zero indicate that the MoDT model provided a better fit.

In this analysis we considered only isotropic matrices ***Q*** = *q**I***, where *q* is a positive scalar and ***I*** is the identity matrix. When *q* = 0, the MoDT model is equivalent to a stationary (non-drifting) mixture model. As we increase *q*, we allow more drift in the model, and initially we find that this improves the quality of fit for all units. If we further increase *q*, we find that the quality of fit eventually diminishes due to overfitting (see appendix C.4 for more detail).

The optimal value of q varies across units (Figure 3C, bottom) and depends on the stability of the tetrode and the firing rate of the unit. Since we use the same value of *q* for all units, we chose a relatively low value (2 μV^2^/hr), which is lower than optimal for many units but still outperforms a stationary model for the vast majority of units. This produces a smoothed estimate that may not follow all of the fluctuations in cluster location, but is still able to capture slower trends (see e.g. Figure 1C). Despite this excessive smoothing, we still find that cluster drift accounts for 12-30% of the spike variability observed in longer recordings (Figure 3D).

### 3.2. Heavy-tailed residuals in empirical data

We also quantified the heavy-tailed distributions of the spike clusters. First, we note that these heavy tails are present even in the extracellular background noise when no spikes are detected (Figure 4A). This heavy-tailed noise could arise from the spiking of many distant neurons, which may exhibit correlated activity that prevents the central limit theorem from applying.

For single units, we found that the Gaussian distribution dramatically underestimates the fraction of spikes that are located away from the cluster center (Figure 4B,C). Again, the observed distribution is more consistent with a *t*-distribution than a Gaussian.

In the MoDT model, the parameter *v* is a user-defined constant that controls the heavy-tailedness of the assumed spike distribution. At *v* = 1, it corresponds to a Cauchy distribution, which has in nite variance. As *v* → ∞, it approches a Gaussian distribution. We found that most units were best fit with *v* in the range 3-20 (Figure 4D), with some differences between brain areas and cell types. For comparison, Shoham et al. (2003) reported a range of 7-15 for single-electrode recordings in macaque motor cortex. We performed spike sorting using *v* = 7 as this provided a good approximation to both limits of the observed range.

### 3.3. Consequences of using a stationary model

Cluster drift is a well-known feature of chronic recordings, and many techniques have been proposed to address this phenomenon. A common approach is to break the recording into chunks, perform spike sorting on each chunk independently, and finally link the clusters across time (Bar-Hillel et al., 2006; Tolias et al., 2007; Wolf & Burdick, 2009; Shalchyan & Farina, 2014; Dhawale et al., 2015).

This approach comes with a tradeoff: short chunks may not contain enough spikes from low-firing neurons, but long chunks suffer more from the effects of drifting clusters. We characterized this tradeoff by breaking our recordings into chunks of varying duration, re-fitting each chunk with a stationary model, and analyzing the result. We identified three common failure modes of this approach (Figure 5): (A) fragmented units due to non-clusterable chunks, (B) loss of isolation between units, and (C) splitting of single units.

Unit fragmentation occurs when a unit is unable to be linked across chunks. The proposed linking algorithms do not link units over a gap in activity, so a single non-clusterable chunk will break the chain of linked units. We evaluated this by counting how many spikes a given unit fired within each chunk, and we considered any chunk with fewer than 25 spikes to be non-clusterable for that unit (Figure 5A). Figure 5D shows the overall fraction of non-clusterable chunks (dashed green line) and the fraction of units that are thus fragmented (solid green line).

Longer chunks are therefore needed to ensure that each chunk contains enough spikes to prevent unit fragmentation. However, longer chunks expose us to more cluster drift, which can cause a loss of isolation and splitting of single units.

Loss of isolation occurs when two drifting clusters occupy the same region of feature space at different times, which appears as cluster overlap under a stationary analysis (Figure 5B). Figure 5D (line B) shows the fraction of our well-isolated units would have failed to meet our quality threshold if had instead used a stationary model to evaluate the unit isolation.

Cluster drift may also produce a multi-modal distribution that leads to a single unit being split into two clusters (Figure 5C). We quantified this effect by identifying cases where the Bayes information criterion (BIC) would justify splitting a cluster into two (Figure 5D, line C1). In some cases, the resulting clusters are well-isolated from one another (less than 5% overlap; Figure 5D, line C2) and would likely require timing information to identify them as a spuriously split unit.

These tradeoffs are faced by any approach, whether model-based or not, that performs spike sorting on each chunk independently. Although the MoDT model also uses discrete time frames, it avoids this tradeoff by aggregating data across frames: it uses the same cluster scale matrix ***C*** for all time frames and incorporates a drift regularizer that effectively smoothes the estimated cluster location ***μ*** over time. As a result, it is able to track units regardless of how few spikes it may fire in a given time frame, which enables us to use sufficiently short time frames (1 minute) that the effects of drift are negligible.

### 3.4. Sensitivity analysis of unit isolation metrics

Spike sorting produces labeled clusters of spikes, also known as units. The spike trains of well-isolated units are typically interpreted as the spiking activity of individual neurons. However, spike detection and sorting are not perfect, and a given unit may contain spurious spikes (false positives) or may fail to capture all spikes from a given neuron (false negatives). Depending on the scientific question being addressed, our subsequent analysis may be more or less sensitive to the presence of such errors. Reliable, quantitative measures of unit quality are therefore critically important for the proper interpretation of the spike sorting output.

Isolation distance and L-ratio (Schmitzer-Torbert et al., 2005) are two such metrics that have found widespread use. Below we consider how well these measures compare to estimates of misclassification error derived from a Gaussian model (Hill et al., 2011), K-means consensus (Fournier et al., 2016), and the MoDT model we propose.

Isolation distance and L-ratio are based on the Mahalanobis distance *δ_nk_* from cluster *k* to spike *n*. If there are *N_k_* spikes in cluster *k*, then its isolation distance is the *N_k_*th smallest value of *δ_nk_*^2^ among the spikes not assigned to that cluster. L-ratio is defined as *L/N_k_*, where *L* is the sum, over all spikes *n* not assigned to cluster *k*, of the complementary CDF of a 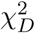 distribution evaluated at *δ_nk_*^2^. This summand can be interpreted as the P-value, using the Mahalanobis distance as the test statistic, under the null hypothesis that the given spike came from a Gaussian distribution fitted to the spikes assigned to cluster *k*.

Misclassification errors are spikes assigned to a cluster that should have been assigned to a different cluster. These may be estimated from a generative model as we describe in section 2.6; we tested both a Gaussian distribution and a *t*-distribution with *v* = 7. K-means consensus is a non-model-based approach in which K-means is used to partition the data based on their Euclidean distance in feature space. This is repeated multiple times from random initializations, and the estimated misclassification error is computed from the fraction of a given cluster’s spikes that have been co-partitioned with other clusters’ spikes.

How do these metrics handle the variations in cluster size, scale, and tail distribution that we observe in our empirical data? To answer this, we first measured the relative size (*N_k_*) and scale (trace(*C_k_*)) of the nearest neighboring cluster for each of our well-isolated units (Figure 6A). This shows that potential sources of contamination may come in a wide variety of sizes and scales. We chose 10 representative cases and synthesized noise clusters of those sizes and scales (Figure 6B). We used that synthetic data to compare the output of these unit isolation metrics with the true overlap between clusters (Figure 6C). By repeating this simulation with clusters of different sizes and scales, we can analyze the sensitivity of these metrics to these extraneous factors (Figure 6D). We synthesized these clusters using a *t*-distribution (*v* = 5:5) in a 12-dimensional feature space.

We also evaluated each of the five metrics on our empirical data (Figure 6E). To create a basis for comparison, we created an aggregate measure of unit isolation based on all five metrics combined (see appendix C.5 for more detail). A broad distribution (e.g. isolation distance) indicates a metric that disagrees with the others about the relative isolation quality of many clusters. We use this empirical comparison and the simulated sensitivity analysis to evaluate the isolation metrics under consideration.

Isolation distance suffers from one important flaw: the contaminating cluster is completely ignored if it contains fewer spikes than the cluster being measured. In such cases, the isolation distance is determined by the location of the second-nearest cluster, and may be arbitrarily large. As a result, a large isolation distance does not necessarily imply good isolation, particularly for units with many spikes.

L-ratio is a more informative metric, but its value can be difficult to interpret. The relationship between the L-ratio and the true error rate is highly dependent on the dimensionality of the feature space and the heavy-tailedness of the distribution (Figure B.1). This makes it difficult to compare L-ratio thresholds across experimental settings unless the underlying noise statistics are known.

**Figure B.1.**
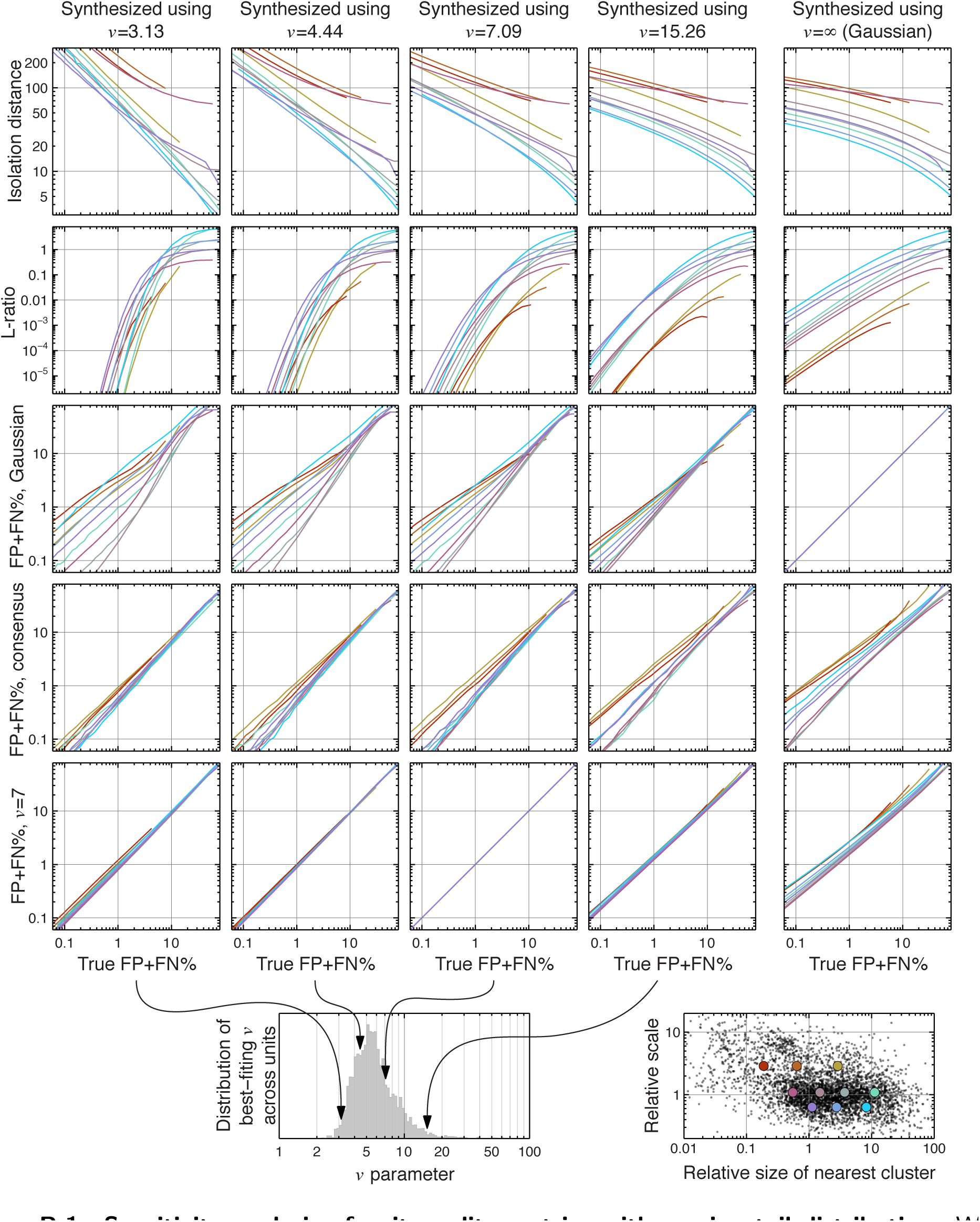
Sensitivity analysis of unit quality metrics with varying tail distribution. We repeated our sensitivity analysis (Figure 6C) using a range of *v* for the synthetic clusters. The *v* parameter controls the heavy-tailedness of the *t*-distribution, with smaller values corresponding to heavier tails. We chose four values of *v* based on the distribution we observed in our well-isolated units (bottom left), and also included a Gaussian case for reference. Note that the values of isolation distance and L-ratio vary dramatically over this range of *v*. The estimated FP+FN using a Gaussian model provides exact results when the data are Gaussian, but becomes sensitive to the relative size/scale of the contaminating cluster when the data come from a heavy-tailed distribution. In contrast, the estimated FP+FN using a *t*-distribution model remains insensitive to cluster size/scale and provides a relatively accurate estimate across the range of tested parameters.

The remaining metrics—total FP+FN based on a Gaussian model, K-means consensus, or a *v* = 7 *t*-distribution model—achieve fairly similar performance when the misclassification error is large. The Gaussian model’s estimates begin to diverge for very well-isolated units, but such inaccuracies may be irrelevant if such units already meet the necessary criteria on unit isolation quality.

K-means consensus exhibits very good performance on the synthetic clusters. However, we encountered a few issues when applying it to empirical data. First, we had to break long datasets into chunks in order to account for drift, and were thus faced with the tradeoff analyzed in Figure 5. During periods when a unit is silent, it is assigned zero false negatives and it cannot contribute to other units’ false positive counts. Second, we encountered unreliable estimates for units that contributed a small fraction of the overall spikes. In such cases, the lack of a consistent K-means partitioning was due to the small number of spikes in the cluster rather than overlap with neighboring clusters. Finally, this metric took much longer to compute than the other metrics tested.

Finally, the *t*-distribution model performs nearly perfectly on the synthetic clusters, but this is not surprising since these data were synthesized using a similar distribution. However, we also found that it continues to provide accurate estimates over a wide range of *v*, including the Gaussian case (Figure B.1). Furthermore, it provides accurate estimates of FP and FN separately, which the Gaussian model and K-means consensus were not able to do (Figure B.2).

**Figure B.2.**
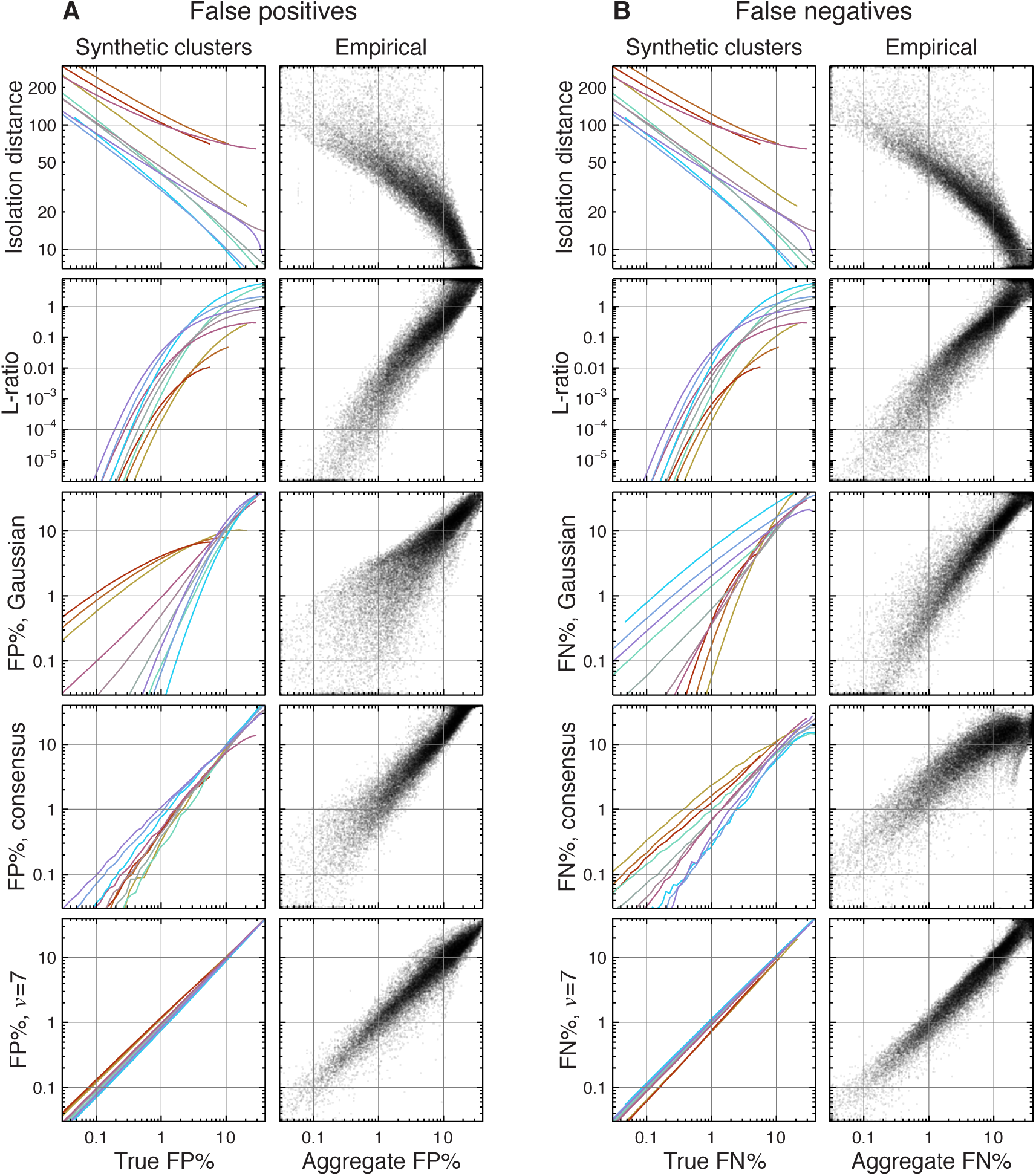
Separate analysis of false positive and false negative estimates. We performed the analysis in Figure 6 separately for the false positive (FP) and false negative (FN) estimates. Note that the Gaussian model and K-means consensus both show increased sensitivity to the size/scale of the contaminating cluster. However, the effects on the FP and FN estimates tended to be opposite in sign for a given case, partially canceling when we consider the sum FP+FN, as in Figure 6.

## 4. Discussion

### 4.1 Spike sorting for chronic recordings

Continuous recordings over the course of days or weeks can be a powerful tool for studying the long-term dynamics of neural firing properties (Harris et al., 2016). This requires us to track single units over long periods of time, and model-based clustering using the MoDT model is well-suited for this task.

In particular, establishing when a neuron is silent is often just as important as knowing when it is firing. However, this involves demonstrating an absence of spikes in the region of feature space where we expected to see them, and this fundamentally requires a model-based approach.

This is why we did not consider the possibility of linking over non-clusterable chunks in our earlier analysis (section 3.3). Although the proposed linking algorithms could be modified to link over a non-clusterable chunk, doing so poses a problem when using the sorted spike trains to draw conclusions about neural activity. Linking a unit over a non-clusterable chunk would imply that it was silent during this period. However, one’s inability to cluster a unit in a given chunk does not certify that it was silent; it could have fired insufficient spikes to warrant its own cluster or it could have been spuriously merged into another cluster.

In contrast, the MoDT model effectively interpolates the cluster’s expected location between consecutive “sightings” of the unit, giving us a reasonable guarantee that the lack of spikes assigned to this unit in the intervening period is indeed due to its silence. Although linking is still necessary for continuous recordings, the MoDT model simplifies the linking process by allowing us to perform spike sorting in segments up to 10 hours in duration. The use of longer segments ensures that all units will fire enough spikes to be clustered, reduces the number of segments that need to be linked, and enables the use of overlapping segments. For example, a week-long recording can be broken into 21 ten-hour segments with an overlap of 2 hours each. We can then establish cluster correspondences based on the spike assignments of the overlapping data.

### 4.2. Measuring unit isolation quality in chronic recordings

The analysis of long recordings also requires unit quality metrics that can handle drift. The MoDT model accomplishes this by explicitly tracking the clusters over time. The use of a *t*-distribution also provides a natural robustness to outliers (Figure 2) and produces accurate estimates of misclassification error over a wide range of conditions (Figure 6).

However, the MoDT model is still a highly structured model. Each cluster is elliptically symmetric with a predetermined tail distribution, and the drift regularization discourages sudden changes in the cluster’s location. It is only through slow drift over time that we can trace out an irregularly-shaped cluster in feature space (e.g. Figure 5B). In contrast, non-parametric approaches allow clusters to take on arbitrary shapes, which may require additional review to ensure that they correspond to biophysically plausible spike distributions.

Furthermore, it is important to acknowledge that unit isolation is a time-varying quantity. Drift may cause two clusters to be well-separated at one point in time, but begin to overlap later. Additionally, certain neural phenomena may be associated with increased misclassification error (e.g. synchronous firing of pyramidal cells during hippocampal sharp-wave/ripple complexes). If subsequent analyses are restricted to a particular subset of the overall recording, then the unit isolation measures should be based on those epochs as well. Model-based approaches accommodate this requirement by providing a continuous estimate of misclassification error, which may then be integrated over the appropriate epochs.

Finally, we would like to caution that isolation quality is only one aspect of unit quality overall. Hill et al. (2011) describe a number of additional quality measures. For example, estimating false negatives due to spike detection is an equally important yet frequently overlooked metric. This is especially important in the presence of cluster drift, as fluctuations in spike amplitude may affect detection efficiency, which could manifest as apparent changes in firing rate.

### 4.3. Model extensions

The MoDT model we have presented consists of three components: a spike distribution model, a drift regularizer, and an EM fitting algorithm. These components may be modified or extended in several ways.

For example, we modeled the spike distribution using a *t*-distribution, which is elliptically symmetric. However, some neurons fire bursts in which subsequent spikes exhibit a reduced amplitude, producing a skewed distribution that has a longer tail in one direction. This one-dimensional skew may be modeled using a restricted multivariate *t*-distribution, which can be fitted using an EM algorithm (Lee & McLachlan, 2014).

We also used a very simple form of drift regularization, but this could be replaced with a more sophisticated model. High-density probes may benefit from a model that explicitly accounts for correlated changes in cluster location due to physical motion of a rigid multi-site probe. This would improve tracking of neurons with a low firing rate.

Finally, the EM algorithm has been widely studied and improved upon in many ways. The basic algorithm is is well-suited for large-scale data processing and is amenable to parallel computing on GPU hardware (Table 2) or distributed computing using high-level data flow engines (Meng et al., 2016). A variety of algorithmic approximations have also been developed to improve performance on high-dimensional data, such as masked EM (Kadir et al., 2014), partial E-steps (Neal & Hinton, 1998), and low-rank covariance decomposition (Magdon-Ismail & Purnell, 2010).

The MoDT model thus offers a modular framework that may be readily adapted to experimental needs.

## 5. Conclusion

In this paper we have described a mixture of drifting *t*-distributions (MoDT) model, which captures two important features of experimental data—cluster drift and heavy tails—and is robust to outliers. When used for spike sorting, the MoDT model can increase unit yield by separating clusters that appear to overlap and decrease user workload by reducing the incidence of clusters that are spuriously split due to drift. As a unit isolation metric, this model provides accurate estimates of misclassification error over a wide range of conditions. These features, along with a computationally efficient EM algorithm, make this well-suited for analysis of long datasets.

## Acknowledgments

We thank all the lab members who contributed user-hours towards data collection and spike sorting (C Wierzynski, A Hoenselaar, M Papadopoulou, B Sauerbrei). Special thanks to Alex Ecker for development of the **moksm** MATLAB package, and Andreas Hoenselaar for development of the clustering GUI.

This work was supported by NSF grants 1546280 and 1146871, NIH grant 1DP1OD008255/5DP1MH099907, the Moore Foundation, and the Mathers Foundation.

### A. Additional comments on the M-step *μ* update

In this appendix we discuss the optimization of ***μ*** performed by equation (6) and compare it to other techniques.

First, note that the optimization of ***μ*** depends on the value of ***C*** and vice-versa. Although standard EM algorithm calls for maximizing 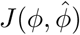 over all *ϕ*, the convergence of a generalized EM algorithm requires only that we improve upon the previous 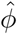 (Dempster et al., 1977). Therefore we need not simultaneously optimize ***μ*** and ***C***, but may instead optimize them one at a time.

The optimization ***μ*** of could also be performed using a Kalman filter with a Rauch—Tung—Striebel backwards pass (Calabrese & Paninski, 2011). Since we have multiple observations per time frame, the forward pass can be performed more efficiently using an alternative parameterization of the Kalman filter known as the *information filter* (see e.g. Anderson & Moore, 1979). In fact, our ***M**_kt_* and ***b**_kt_* correspond to the information matrix and vector, respectively, of the information filter’s observation update step. Nonetheless, we have found that it is faster to solve equation (6) directly using standard numerical linear algebra routines (i.e. LAPACK dpbsv).

For more insight into this optimization, consider the unregularized case (***Q*** → ∞ and hence ***Q*** ^−1^ = 0). In this scenario, the time frames are indepenent of one another (the ***A*** matrix is block diagonal) and the solution to equation (6) is simply

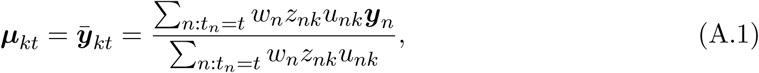

i.e. the weighted sample mean of spikes assigned to cluster *k* in time frame *t*.

As we add regularization, the optimal ***μ*** becomes a temporally smoothed version of this weighted average. Consider for example the case where ***Q*** and ***C***_k_ are isotropic, i.e. ***Q*** = q***I*** and ***C***_k_ = c***I***, where *I* is the identity matrix. In this scenario, we can rewrite equation (6) as

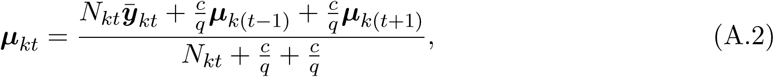

where 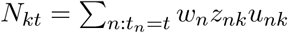 is the weighted number of spikes assigned to this cluster in this time frame. Equation (A.2) is analogous to a bi-directional exponentially-weighted moving average. Each ***μ**_kt_* is a weighted average of that time frame’s sample mean 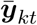 and its neighbors ***μ***_k(t-1)_ and ***μ***_k(t+1)_.

In time frames with no spikes (*N_kt_* = 0), ***μ**_kt_* simply takes the midpoint between its neighbors. As we increase the number of spikes (*N_kt_* ↑), ***μ**_kt_* places more weight on its own sample mean rather than interpolating between its neighbors. This is evident when comparing the high-and low-firing units in Figure 1C. Likewise, tightening the cluster variance (*c* ↓) or increasing the expected drift (*q* ↑) will also shift the average in equation (A.2) in favor of the sample mean.

Finally, note that the equations in section 2 and in this appendix use units of [feature space units]^2^/[time frame] for ***Q*** = *q**I***. In contrast, Figure 3 and section 3.1 report *q* in units of [feature space units]^2^/hr for ease of interpretation.

### B. Additional analysis of unit isolation metrics

In this appendix we perform additional analysis of unit isolation metrics to supplement Figure 6.

Figure B.1 repeats the sensitivity analysis of Figure 6D using a range of distributions for the synthetic clusters. As we vary the heaviness of the distribution tails, we find that relationship between the true error rate and the isolation distance or L-ratio can vary dramatically. For example, an L-ratio of 0:01 corresponds to an FP+FN between 1.5-6% for a heavy-tailed distribution (*v* = 3.1), but could range anywhere from 0.1-10% for a Gaussian distribution. These values of these metrics also depend strongly on the dimensionality of the feature space. In contrast, the *v* = 7 model-based estimate (and the consensus-based estimate, to a lesser degree) remains insensitive to the cluster size/scale and provides an accurate estimate of misclassification error across the range of distributions.

Figure B.2 repeats the sensitivity analysis (Figure 6D) and empirical comparison (Figure 6E) for the FP and FN separately. Here we see that the Gaussian-and consensus-based estimates exhibit increased variability and bias when asked to estimate FP and FN separately, whereas the *v* = 7 model continues to provide accurate results.

### C. Analysis methods

This appendix contains additional details on the analyses performed to generate the figures in this paper.

#### C.1. Theoretical distribution of Mahalanobis distance

Figure 1D shows the distribution of the (non-squared) Mahalanobis distance *x* computed using the fitted ***μ*** and the sample covariance Σ

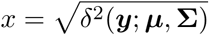

If *y* come from a multivariate Gaussian distribution and we assume that ***μ*** and **Σ** are accurate estimates of the true mean and covariance, then *x*^2^ will be distributed according to a chi-squared distribution with *D* degrees of freedom. We can apply a change of variables to obtain the theoretical probability distribution function (PDF) of the non-squared *x*:

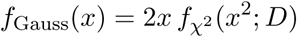

where 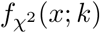 denotes the PDF of a chi-squared distribution with *k* degrees of freedom. This *f*_Gauss_(*x*) is the dashed line in Figure 1D, and its corresponding cumulative distribution function (CDF) is shown in Figure 4B, C.

If *y* come from a multivariate *t*-distribution with *v* degrees of freedom, then the quantity (1/D) *δ*^2^(***y***;***μ***, ***C***) will be distributed according to an F distribution with (D, *v*) degrees of freedom (Box & Tiao, 1973). However, our Mahalanobis distance *x* is computed using the sample covariance Σ rather than the *t*-distribution scale parameter C. If we assume that Σ equals the expected covariance (this requires *v* > 2, as the expected covariance is undefined or infinite when *v* ≤ 2), then δ^2^(***y***;***μ***, ***C***) = *vx*^2^/(*v*-2), and applying this change of variables gives us 
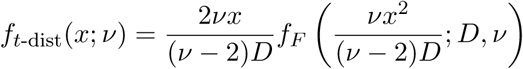
where *f*_F_ (*x*; *k*_1_, *k*_2_) denotes the PDF of an F distribution with (*k*_1_, *k*_2_) degrees of freedom. This *f*_t-dist_(*x*;*v*) is the solid line in Figure 1D, and its corresponding CDF is shown in Figure 4C.

#### C.2. Robust covariance estimation

In this paper we have focused on data with heavy tails (Figure 4), but the *t*-distribution’s robustness to outliers suggests that it may be a useful model even when the data are Gaussian (Figure 2).

However, fitting a *t*-distribution to Gaussian data causes us to overestimate the distribution tails. In Figure 2A, note how the fitted *t*-distribution’s 99% confidence ellipse (green) is inflated relative to the true ellipse (black). In this situation, we would like to use the fitted *t*-distribution to derive robust estimates of the Gaussian parameters.

As the number of samples *N* → ∞, the fitted ***μ*** converges to the true mean of the Gaussian distribution. However, the fitted ***C*** is a biased estimator of the true covariance ***Σ***. If we assume the data are Gaussian, we can compute a correction factor by solving β for in the following:

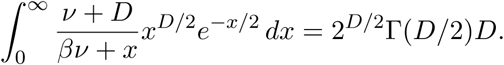

We can then use *β**C*** as a robust estimate for the Gaussian covariance. The corresponding 99% confidence ellipse is actually shown in Figure 2A (light grey ellipse in right panel), but it is visually obscured by the 99% confidence ellipse of the true distribution (black).

#### C.3. Relative influence of single spikes on fitted parameters

In Figure 2B we analyze the relative influence of a single spike on the fitted parameters. This appendix describes how this "relative influence" is computed.

The top panel shows the relative influence on the fitted ***μ*** as a function of the spike’s distance from the cluster center. To compute this quantity, let 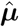 denote the fitted cluster location with spikes, and consider the effect of adding a spike *y*_N+1_. In the case of a single cluster and a single time frame, the M-step update (equation 6) gives us

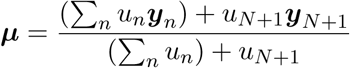

If the new spike is located at a distance 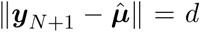 from the cluster center, then the change in the fitted ***μ*** is

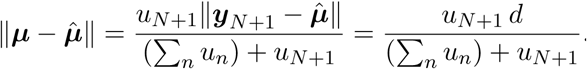

We will denote this quantity *I_μ_*(*d*), the influence of a single spike at a distance *d*. For simplicity, let us assume that *N* is large (and hence 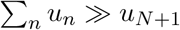) and that the cluster scale ***C*** = c***I***.

Under these assumptions,

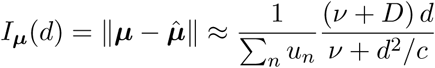

Note that *I_μ_*(*d*) → 0 as *d* → ∞.

In Figure 2B we consider the case where the true data are distributed according to a standard multivariate normal distribution (Σ = I). For a given *v*, the fitted cluster scale can be determined using the procedure in appendix C.2. The top panel of Figure 2B shows the relative influence 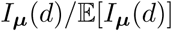, where 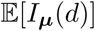 is the expected value of *I_μ_* over the spike distribution.

The bottom panel of Figure 2B repeats this analysis for the scale parameter ***C***. Again assuming that ***C*** = *c**I***,

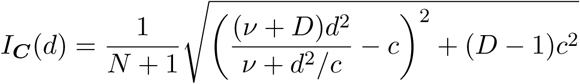

#### C.4. Model assessment using the likelihood ratio

Figures 3C and 4D use the log likelihood ratio (LLR) to evaluate the quality of fit of a MoDT model under various choices for the user-defined model parameters.

The LLR is the logarithm of the likelihood ratio comparing the MoDT model to some alternative (a stationary t distribution in Figure 3 and a drifting Gaussian distribution in Figure 4). Values greater than zero indicate that the MoDT model produced a better t. We report the LLR divided by the number of spikes so that we may compare across datasets of different sizes.

When evaluating the effect of varying the drift regularization parameter ***Q***, it is important to distinguish between fitting the underlying cluster drift and capturing the random noise of the observed spikes. Therefore we performed cross-validation using a holdout, and Figure 3C reports the LLR evaluated on the validation test set. This was performed by randomly partitioning the spikes into two equal-sized subsets (a training set and a test set) and fitting the models to the training set only. Note that relaxing the drift regularization (i.e. increasing ***Q***) will always improve the model’s ability to t the training data. However, the likelihood of the test set increases initially (due to the model’s improved ability to track the cluster drift) but eventually decreases due to over tting to the training set.

#### C.5. Aggregation of isolation metrics

In order to compare the output of the five unit isolation metrics we tested, we constructed an aggregate measure based on the five metrics combined (Figure 6E).

Since two of these metrics use a different scale than the rest, we first constructed a relative ranking of clusters for each metric separately. We then combined these five rankings into an aggregate ranking using Borda’s method followed by local Kemenization (Dwork et al., 2001). This procedure ensures that any cluster *i* that is considered better-isolated than cluster *j* by a majority of metrics will be ranked higher in the aggregate ranking.

With the clusters thus ranked, we assigned each cluster an aggregate FP+FN value based on the cumulative distribution of reported FP+FN from the three metrics that report in this scale. This preserves the relative ordering defined by the aggregate ranking.

